# Personalized intrinsic network topography mapping and functional connectivity deficits in Autism Spectrum Disorder

**DOI:** 10.1101/161893

**Authors:** Erin W. Dickie, Stephanie H. Ameis, Joseph D. Viviano, Dawn E. Smith, Navona Calarco, Saba Shahab, Aristotle N. Voineskos

## Abstract

**Background:** Recent advances demonstrate individually specific variation in brain architecture in healthy individuals using fMRI data. To our knowledge, the effects of individually specific variation in complex brain disorders have not been previously reported.

**Methods:** We developed a novel approach (Personalized Intrinsic Network topography, PINT) for localizing individually specific resting state networks using conventional resting state fMRI scans. Using cross-sectional data from participants with ASD (n=393) and TD controls (n=496) across 15 sites we tested: 1) effect of diagnosis and age on the variability of intrinsic network locations and 2) whether prior findings of functional connectivity differences in ASD as compared to TD remain after PINT application.

**Results:** We found greater variability in the spatial locations of resting state networks within individuals with ASD as compared to TD. In TD participants, variability decreased from childhood into adulthood, and increased again in late-life, following a ‘U-shaped’ pattern, which was not present in those with ASD. Comparison of intrinsic connectivity between groups revealed that the application of PINT decreased the number of hypo-connected regions in ASD.

**Conclusions:** Our results provide a new framework for measuring altered brain functioning in neurodevelopmental disorders that may have implications for tracking developmental course, phenotypic heterogeneity, and ultimately treatment response. We underscore the importance of accounting for individual variation in the study of complex brain disorders.

## Introducton

Disease heterogeneity has been a major obstacle in the study of neurodevelopmental disorders. Autism spectrum disorder (ASD) is a lifelong complex neurodevelopmental disorder affecting 1% of the population(1), and has been characterized by both ‘hyper-connectivity’, and ‘hypo-connectivity’, using neuroimaging, neuropathology, and other investigative techniques. Such heterogeneity is often attributed to different disease subtypes, IQ, MRI scanner differences, age range, sex, or treatment effects. However, variability among individuals is a topic of emerging interest that may have broad implications for our understanding of neurodevelopmental disorders, such as ASD, as well as ultimately in guiding approaches to treatment.

Each human being may possess his or her own unique intrinsic neural topography(2). Individually specific topography can be measured reproducibly in healthy individuals using intrinsic connectivity(3–6). At the individual level, Hahamy and colleagues(7) found greater spatial variability of inter-hemispheric connectivity in approximately 80 ASD cases compared to 80 controls, raising the question of whether such increased variability exists within specific brain networks. Further, it raises several questions: a) is such individually specific topography different in a complex neurodevelopmental disorder such as ASD compared to TD controls, b) how does individually specific topography change in atypical versus typical development, i.e., with age? and c) what does the presence of individually specific topography mean for interpretation of group-based findings in ASD and other neurodevelopmental disorders that are reported in the literature?

To address these questions, we developed an iterative algorithm, Personalized Intrinsic Network Topography (PINT) and made it publicly available. For each participant in a dataset, the PINT algorithm shifts the locations of so-called “template” ROIs to a nearby cortical location, or “personalized” ROI, that maximizes the correlation of the ROI with the rest of the ROIs from its network. Here, we applied the PINT algorithm to resting state fMRI data in a large sample from the ABIDE I dataset(8). We first tested the stability/reliability of intrinsic network identification using PINT in the same participants over time with the longitudinal subsample of the ABIDE dataset (n=31, from two sites). We then proceeded to test the following main objectives using ABIDE cross-sectional data from participants with ASD (n=393) and TD controls (n=496) across 15 sites: a) is variability of intrinsic networks increased in ASD compared to TD controls, thus acting as a marker of ASD; b) what is the effect of age (and potentially neurodevelopment) on intrinsic network variability, and; c) do prior findings of functional connectivity differences in ASD versus TD remain after PINT application.

## Materials and Methods

### Datasets

We employed the publicly available ABIDE I resting state dataset (8) of ASD and TD participants from 17 sites. For details of the scanning parameters for each individual site see http://fcon_1000.projects.nitrc.org/indi/abide/abide_I.html. Data from the OHSU site were not included because the shorter duration of the resting state scans was not suitable for FSL’s ICA FIX de-noising (see preprocessing methods). Data from the Stanford site were excluded due to poor quality of the anatomical images leading to a poor performance of our FreeSurfer pipeline. Demographic criteria from participants included in our analyses (n=393 ASD, n=496 TD, across 15 sites, ages 6-65) are described in Supplemental Table S2.

The longitudinal stability of the PINT algorithm was tested in the ABIDE Longitudinal sample, a release of two year follow-up scans from the UCLA (n=21, ages 10-13) and the University of Pittsburgh, School of Medicine (UPSM, n=17, ages 9-17)). Scanning parameters are summarized in Supplemental Appendix 3.

### Preprocessing Pipeline

MR images were preprocessed using a workflow, described in the ‘Preprocessing Pipeline’ section of Supplemental Appendix 1, adapted from the HCP Minimal Processing Pipeline (9) to register data into a combined volume and cortical surface-based analysis format (cifti format in “MNINonLinear-fsaverage_LR32” space). Adapted scripts are available at https://github.com/edickie/ciftify). Briefly, the cortical surfaces were defined using FreeSurfer’s recon-all pipeline (version 5.3). Resting state scans were preprocessed using a combination of AFNI and FSL tools for slice timing, motion correction, and ICA based data-cleaning using ICA FIX.

### The PINT algorithm and template ROIs

Eighty “template” vertices were chosen to sample from six resting state networks described in Yeo et al(10) (Dorsal Attention (DA), Default Mode (DM), Ventral Attention (VA), Fronto-Parietal (FP), Sensory Motor (SM), and Visual (VI)). The “Limbic” Network from the seven network atlas was not employed because it contains areas of high fMRI signal susceptibility. The locations of the 80 ROIs are given in Supplemental Table S1 and plotted in Figure 1. Average seed correlation maps, shown in Supplemental Figures S1-S2, strongly resemble the Yeo 7 network atlas. Therefore, we are confident that these 80 template ROIs represent a good sample of network activity. PINT fits an individual participant's resting connectivity matrix to a “template” pattern of networks by moving the locations for sampling ROIs, iteratively, in a manner that optimizes the within-network connectivity. The code is available at https://github.com/edickie/ciftify, see ciftify-PINT-vertices). During each iteration, for each ROI, PINT calculates the partial correlation of each vertex within a 6mm search radius of a start vertex, and the other ROIs from the same network, and then moves the ROI’s position to the vertex of highest partial correlation (see depiction in Figure 1). Details of the algorithms performance in the ABIDE I sample are given in the Supplemental Appendix 1.

**Figure 1.**
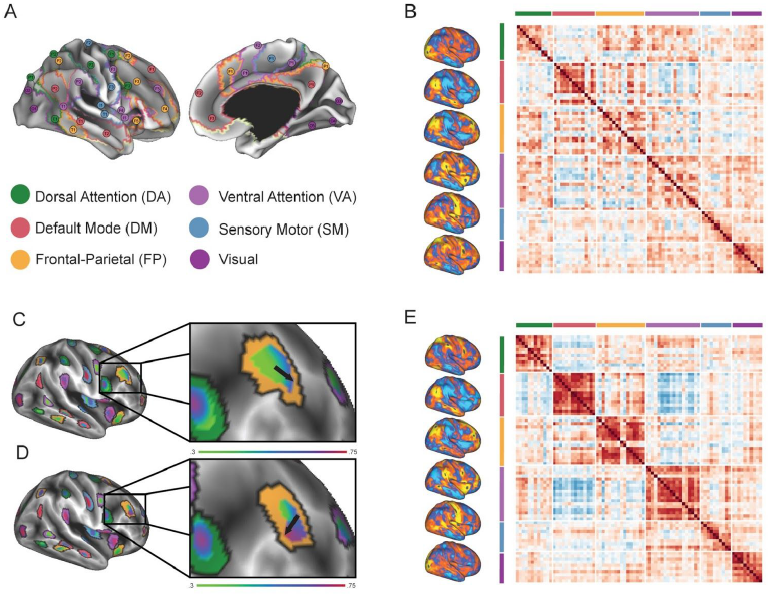
Schematic of Personalized Intrinsic Network Topography (PINT) Method. A) PINT starts with a template set of regions of interest (ROIs) selected from the Yeo et al. (10) atlas. B) Template (input) and E)Personalized (output) correlation matrixes from a representative subject. C) PINT starts by calculating average mean timeseries from circular ROIs (of 6mm radius) around 80 central “template” vertices. Then, for each template vertex, a search area is defined as those vertices within 6mm of the central template vertex and not within 12mm of any other ROI’s template vertex. PINT then calculates the partial correlation of all vertices within a search area around the template vertex and average timeseries from the other ROIs of the same network (the partial correlation controls for the five average timeseries of the ROIs of the other networks) and the center of the ROI is moved to the vertex of maximal partial correlation. D) Once all 80 ROIs have been moved, the algorithm iteratively searches around the new vertex locations. Abbreviations: DA = Dorsal Attention Network, DM = Default Mode Network, FP =Fronto-Parietal Network, VA = Ventral Attention Network, SM = Sensory Motor Network, VI = Visual Network. (Note: the Limbic Network was not included because it contains many areas of high fMRI signal susceptibility.)

### Statistical Analyses

Longitudinal stability was measured by comparing the distance (measured on the HCP S900 average surface mid-surface) between the baseline and follow-up personalized ROI locations. Within participant distance was compared to the mean cross-subjects distance with paired t-tests.

Individual variability in intrinsic network locations was calculated as the distance from the starting “template” vertex location to each participant’s personalized vertex location. This distance was averaged across all 80 ROIs for each participant to build a “total brain” score, and within each network to produce network level scores. We tested for effects of diagnosis, and age-by-diagnosis interactions on average distance using linear regression. All linear models included covariates of age, sex, IQ, site, cortical surface area and the top two principal components of scan quality metrics. To identify non-linear age effects, the “total brain” score was regressed against the same model with a quadratic age term introduced and the same covariates. To confirm the nature of age effects in those 30 and under, both the non-linear (quadratic) and linear models were repeated after excluding participants (37 ASD, 33 TD over the age of 30). Network level analyses were corrected for multiple comparisons using false discovery rate (FDR).

To visualize personalized ROI locations within groups of people, we created vertex-wise probability maps. A difference map was created by subtracting the probability map of the ASD participants from that of TD participants. This map was thresholded using permutation tests. We permuted group membership in order to created 2000 “random” group difference maps. Then, at each vertex, we took difference values between the 2.5th and 97.5th centiles as our cut-off for significance (alpha 0.05, two-tailed). A similar process was used to create difference maps between children and young adults.

Edges from Z-transformed correlation matrices calculated from timeseries of: 1) pre-PINT adjustment “template” ROIs, and 2) post-PINT “personalized” ROIs were tested for a difference between ASD and TD groups, in a linear regression with covariates of age, sex, IQ, scanning site, and the two top principal components of scan quality measures, using linear regression. FDR was used to correct for multiple comparisons. Follow-up analyses, testing for associations of edgewise connectivity with Autism Diagnostic Observation Schedule (ADOS) scores and Full IQ were conducted using those participants with ASD who had data for these scales using the same covariates (age, sex, site and the top two principal components of scan quality measures).

## Results

### Longitudinal Reliability/Stability of the PINT algorithm

We found that the within-subject distance was lower than the cross-subject distance both imaging sites: UPSM (t(16)=8.28, p=3.5e-07) and UCLA (t(13)=5.99, p=4.5e-05)(see Figure 2). This effect was significant for all six resting state networks tested (see Supplemental Figure S7 and Supplemental Table S5), and irrespective of ASD or TD group membership (see Supplemental Table S6). It should be noted that the sample from UPSM and UCLA was developmental (ages 9-17); thus some re-organization of intrinsic connectivity is expected between timepoints,(11, 12). We also found that greater within-subject than cross subject similarity of the resulting correlation matrixes (described in Supplemental Appendix 2 and Supplemental Figure S8).

**Figure 2.**
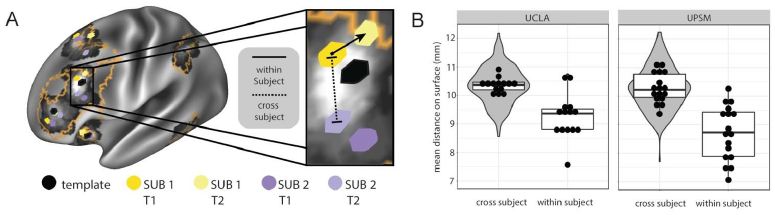
Stability of PINT results in ABIDE Longitudinal Dataset. A) Measurement of within-subject and cross-subject distance for a personalized Fronto-Parietal Network (FP) ROI is depicted for two representative longitudinal participants. The orange outline represents the Fronto-parietal Network as defined by the Yeo et al. (10) atlas. “Template” vertex locations are shown in black for reference. b) The locations of PINT “personalized” vertices show consistency over time. Grey violin plots show the distribution of cross-subject distances (averaging across 80 ROIs) for the UCLA site (left, n=14) and UPSM site (right, n=17). Box plot and points, for the cross-subject measure, show the mean distance of each subject’s baseline scan to all other subjects’ follow-up scan. This measure is compared with the distance of each subject’s baseline scan to their own follow-up scan (within subject distance).

### Intrinsic Network Locations are more variable in ASD participants compared to TD participants

The personalized ROI centers were within an average of 7.70mm of the template vertex locations (see Supplemental Figure S3 for more detailed probabilistic maps). We found that individual subject distances to the template ROI center were greater in the ASD group compared to the TD group (t(867)=3.05, p=0.002).

### Intrinsic Network Location Variability follows a U shaped curve pattern across the lifespan in the TD group, but shows no relationship with age in the ASD group

Within the TD group, the mean distance from the template ROI center to personalized ROI center (across the 80 ROIs tested) was negatively associated with age (t(475)=-3.17, p=0.002). Under further exploration, the effect of age in the TD group was best described by a quadratic curve, where the total distance decreased from childhood to adulthood but then increased again after age 30 (see Supplemental Figure S4, quadratic term t(474)=3.07, p=0.002). If only those TD participants aged 30 and below are included, the age effect is best modeled as a linear decline (t(442)=-4.26, p=2.5e-05). ASD participants showed no such age effect: neither a linear decline (t(371)=-1.49, p=0.14), nor quadratic curve (t(371)=0.31, p=0.75) was observed. However, a significant age-by-diagnosis effect was found (F(2,864)=3.19, p=0.04).

### Network level analysis of intrinsic network locations

Given the age effects described above in TD participants, we assessed each of the six functional networks separately in participants under the age of 30 (see Table 1). We found an effect of diagnosis and a diagnosis-by-age interaction in the Ventral Attention and Dorsal Attention networks. (see Table 1, and Figure 3a). For the Default Mode Network (DM), an effect of age was present (see Table 1)). Spatial variability in DM ROIs decreased with age (ASD: r=-0.11, p=0.05; TD: r=-0.10, p=0.03; Figure 3b). There was no effect of ASD diagnosis. However, an exploratory analysis in ASD participants revealed a weak trend for an association between spatial variability in DM ROIs and ASD symptom severity (ADOS scores: t(247)=1.71, p=0.09 uncorrected).

**Table 1.** Effect of age and ASD diagnosis of personalized vertex locations

| Network | Age Effect | Diagnosis Effect | Age by Diagnosis Interaction |
| --- | --- | --- | --- |
| SM | - | $t=-1.75, p=0.080$ | - |
| VI | - | - | $t=-1.89, p=0.060$ |
| DM | $t=-2.87, p<0.001^*$ | - | - |
| FP | - | - | - |
| DA | - | $t=-2.73, p=0.007^*$ | $t=-2.06, p=0.040$ |
| VA | - | $t=-2.76, p=0.006^*$ | $t=-2.02, p=0.040$ |
| All ROIs | - | $t=-3.23, p=0.001$ | $t=-2.49, p=0.013$ |
| *Significant after false discovery rate correction across 6 networks<br>Additional model covariates: Site, Sex, Full IQ, total cortical surface area, top two principal components of fMRI scan quality. Residual degrees of freedom = 886. |  |  |  |

**Figure 3.**
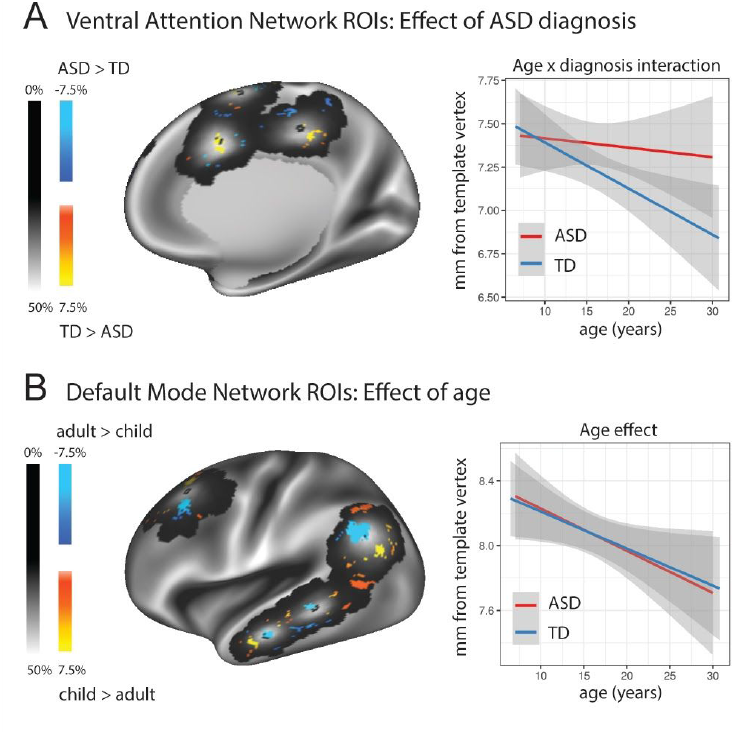
A) Medial right hemisphere view of diagnosis effect on personalized ROIs locations in the Ventral Attention (VA) Network. The probability map of the personalized ROI location (i.e. one where the value at each vertex represents the proportion of participants in the sample, pooling ASD and TD, whose 6mm ROI emcompases that location) is plotted in grayscale. The template vertex locations are plotted as black dots. The overlaid colours show areas where a significant effect of diagnosis was observed (warm colors TD > ASD; cool colours ASD > TD). B) Lateral left hemisphere view of age effect on personalized ROIs locations in the Default Mode (DM) Network. The probability map of the personalized ROI location (all participants 30 and under) is plotted in grayscale. The template vertex locations are plotted as the black dot. The overlaid colours show areas where a significant effect of age group was observed (warm colors children > adults; cool colours adults > children). Children were defined as those under the age of 12 (n=183);adults were defined as between the ages of 18-30 (inclusive, n=251).

**Figure 4.**
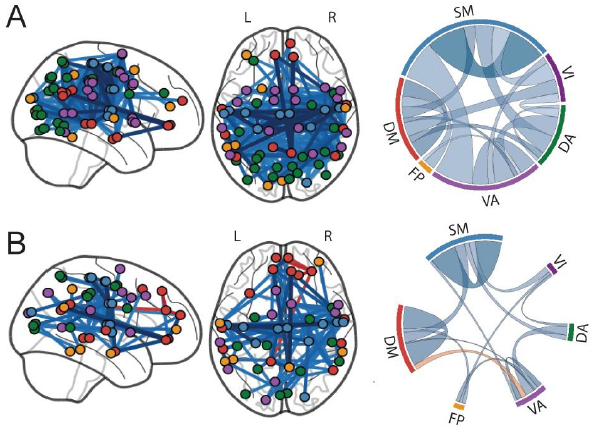
A) Locations (left) and chord diagram of network composition (right) for the 214 edges showing significant hypo-connectivity in ASD calculated from the Template ROIs (before PINT). B) After PINT, the number of significant hypo-connected edges (blue) is reduced 80 and four hyper-connectivity connected (orange) edges are seen. For the chord diagrams, the width of the chord represents the number of significant edges and color of the chord represents the mean effect size for those significant edges.

### Application of PINT (i.e. adjusting individual differences in spatial location) decreases the long range “hypo-connectivity” otherwise observed in ASD

The average (n=889) Z-transformed correlation values are shown in Supplemental Figure S5. Within resting state network functional connectivity increased substantially across the sample of ASD and TD participants when using personalized ROI locations (template ROIs mean(SD) = 0.34(0.12); personalized ROI mean(SD) = 0.52(0.13), see Supplemental Table S3).

Before PINT correction, 214 edges showed significantly lower correlation strength in the ASD group compared to the TD group. This is in high agreement with previous publications of cortical hypo-connectivity in ASD (8). In contrast, after adjusting for differences in intrinsic network topography, 80 connections showed lower correlations in ASD versus TD groups, and four connections showed higher correlation in ASD versus TD groups. This provides direct support for our hypothesis that some of the cortical hypo-connectivity previously reported in ASD is a product of greater heterogeneity in the spatial locations of the ROIs in the ASD versus TD group.

### Effects of clinical scores after PINT application

ADOS total score was negatively correlated with connectivity between the ventromedial prefrontal cortex ROI of the Default Mode Network and the frontal pole ROI of the Ventral Attention Network (ADOS Total Score: t=19.3, p=0.05, FDR corrected). Several additional edges connecting the Ventral Attention and Default Mode Networks showed a similar negative correlation at a lower threshold (p<0.001, uncorrected, see Supplemental Table S4 and Supplemental Figure S6).

## Discussion

We found that the spatial organization of the brain’s intrinsic functional networks is more variable in ASD participants compared to TD participants, most prominently in the Dorsal and Ventral Attention Networks. Application of our novel PINT algorithm to a conventional group-based analysis of functional connectivity resulted in a re-evaluation of many of the previously shown findings of ‘hypo-connectivity’ in the ASD literature. Intrinsic network location variability showed a non-linear U-shaped relationship with age across the lifespan in TD participants, but there was no relationship with age in those with ASD. Our findings provide a new understanding of brain network organization and heterogeneity in people with ASD and TD individuals across the developmental lifespan, while re-configuring our understanding of prior findings in the literature.

Our most important finding was greater variability of spatial organization in cortical resting state networks in the ASD population than in TD controls. Greater heterogeneity of cortical organization in ASD has been reported recently in spatial patterns of homotopic connectivity(7), cortical organization(13), altered functional organization of the motor cortex(14, 15) The increased variability in functional organization observed in ASD aligns with the broad clinical and genetic heterogeneity associated with altered neurodevelopmental trajectories(16). Recent data estimates that between 400-1000 genes are involved in illness susceptibility in ASD(16), with no one genetic polymorphism predicting more than 1% of ASD cases(17). The function of many of these genes appears to converge on molecular pathways for early cortical patterning(18, 19). However, in mouse models, the effects of the top susceptibility genes for autism each have diverse impacts on brain structure(20). Our findings can be interpreted in the context of theories of autism pathology, which posit an early neurodevelopmental insult(19, 21–23). It is possible that, in the presence of an early neuropathological process, the especially plastic developing brain can maintain the function most critically developing at the time, by re-appropriating tissue near to that which has become damaged or disconnected. In the case of ASD, with a wide heterogeneity of genetic(24) and environmental influences(25, 26) at play, various neuropathological factors could manifest in different cortical locations across individuals(21). A direct consequence could be greater heterogeneity of brain topography.

A key implication of our results is that consideration of increased individual variability among ASD participants might be required for future imaging and post-mortem brain analysis. Recently, Uddin et al.(27) found that maps of the ICA component for the Salience Network could be used to classify ASD and TD participants. Thus, disruption of Salience Network function has recently been proposed as a marker of ASD pathology(28). Of the six networks tested in the present study, the Ventral Attention Network (which encompasses the Salience Network) showed the most spatial variability in ASD. When we conducted the conventional functional connectivity comparison between ASD and TD participants, while accounting for the PINT correction, the finding of cortical “hypo-connectivity” with these ROIs in ASD was reduced. It has been previously proposed that some of the hypo-connectivity measured in ASD, was not a “true” drop in connection strength between two ROIs, but rather attributable to greater heterogeneity across subjects in the locations of the functional areas(7). Our findings confirm this contention on a wide scale, i.e., across brain functional networks, although some networks were more susceptible than others.

In contrast to our results from the Dorsal and Ventral Attention Networks, lower within-network connectivity for the Sensory Motor and Default Mode Networks largely persisted as robust potential neural markers of ASD, even after individual differences in network locations were accounted for. Activity within the Sensory Motor Network(8, 14, 15), as well as connectivity between the sensory motor areas and the subcortex(8, 29), have been consistently reported in the ASD literature. Default Mode Network connectivity decreases have also been frequently reported in the ASD literature(8, 30), and have been of great interest due to the proposed role of this network in mentalizing, a core behavioural deficit in ASD(31). Following PINT, four connections between the Ventral Attention Network and the Default Mode Network showed newly significant hyper-connectivity in ASD, compared to TD. Other connections between the Default Mode Network and the Ventral Attention Network were hypo-connected, and negatively correlated with ASD symptom severity. Given the later maturation of the Default Mode Network, it might generally be less susceptible to spatial heterogeneity effects related to ASD, or more complex relationships such as between network connectivity to Attention Network connectivity in ASD, therefore, requires further exploration.

Our findings also revealed an effect of age, whereby in TD participants, we observed that the distance of participants’ personalized ROI locations from the template vertex location decrease from childhood to adulthood. With increasing age, the organization of resting-state networks look more like the template organization in TD participants. In contrast, variability tends to increase in later-life in the TD group, displaying a retrogenesis-like effect. However, the smaller sample size of older adults examined here limits our ability to make strong conclusions in this regard. The age effects we observed are absent in the ASD population, consistent with an altered neurodevelopmental trajectory. The age trajectory that we found in TD participants is similar to those observed for other measures of brain connectivity across development, including white matter volume (11), and white matter microstructure (12, 32, 33).

A recent paper identified 260 functional cortical areas using the multimodal combination of all imaging data from the Human Connectome Project (HCP; including two hours of resting state fMRI (4). We argue that 80 ROIs from six networks is a reasonable resolution for analysis given that, for some sites, as little as five minutes of resting fMRI data is available and some motion during the scan is unavoidable. We are reassured of the validity of this approach given the similarity of results observed across time in our longitudinal sample. This longitudinal retest reliability was observed both for the personalized ROI locations and for the resulting correlation matrices. However, it is possible that ROIs we identify here may actually be multiple subregions belonging to the same one of six larger scale networks, and that these subregions may contribute differently to ASD pathology. The PINT algorithm identifies the center of functional areas of interest. It is possible that some neural markers of ASD pathology may reside, not at the center of functional areas, but in cortical areas in-between these centers. These template locations of resting state function were defined from young adult resting state data(10). Therefore, it is possible that this template organization of resting state networks is most applicable to the young adult age range.

The current ABIDE data release has a relatively small number of female participants. This is expected as more males are affected by ASD. In this analysis we were underpowered to investigate sex or gender-based differences in network organization, which may manifest differently in males versus females with ASD(34, 35). We also were unable to examine effects of comorbidity, due to variability across sites in the nature of the clinical assessment. Additionally, the full-scale IQ range within this population was mostly reflective of individuals with average intelligence (IQ > 80), and, while IQ was included as a covariate in all analyses, we were unable to examine specific associations.

## Conclusion

We show that the identification of personalized functional network organization is possible using conventional resting state fMRI scans from multiple scanners. These results are reliable over time, suggesting that they indicate a "trait-like" marker of brain organization. We provide evidence that the spatial organization of cortical resting state networks is more variable in people affected by ASD. Furthermore, ASD participants did not demonstrate the age-related decreases in variability observed in TD participants. Finally, our results indicated that many findings of widespread hypo-connectivity in ASD are lost following application of PINT, while other new findings emerge. Taken together, these results underscore the importance of accounting for individual variability in the study of complex brain disorders, and provide a window into neurobiological heterogeneity among individuals that may be relevant for treatment innovation.

## Acknowledgements

ANV receives funding from the CIHR, NIMH, BBRF, CFI, Ontario MRI, and the CAMH Foundation. We would like to thank Martin Lindquist for his guidance in how to measure reproducibility, and Adriana Di Martino for advice about the ABIDE dataset and Dayton Miranda for his support with preprocessing quality assurance. We also thank all sites and investigators who have worked to share their data through ABIDE.

Support for ABIDE coordination and data aggregation was partially provided by NIMH (K23MH087770, R03MH096321, and BRAINSRO1MH094639-01), the Leon Levy Foundation, and by gifts from Joseph P. Healy, the Stavros Niarchos Foundation. Support for ABIDE dataset data collection at each site was provided by NIH (DC011095, MH084164, K01MH092288-Stanford; HD55748, KO1MH081191, MH67924-Pitt; K08MH092697, P50MH60450, R01NS34783, R01MH080826, T32DC008553-USM; K23MH087770, R01HD065282, R01MH081218, R21MH084126-NYU; MH066496, R21MH079871, U19HD035482-UM1&UM2; R00MH091238, R01MH086654, R01MH096773-OHSU; R01MH081023-SDSU; 1R01HD06528001-UCLA1&UCLA2; K01MH071284-Yale; R01MH080721-Caltech), Autism Speaks (KKI, NYU, Olin, UM1&2, Pitt, USM, Yale), NINDS (R01NS048527; KKI), NICHD (Yale, UCLA1&2, CMU), the Simons Foundation (OHSU, Yale, Caltech, CMU), the Belgian Interuniversity Attraction Poles Grant (P6/29;Leuven1&2), Ben B. and Iris M. Margolis Foundation (USM), European Commission, Marie Curie Excellence Grant (MEXT-CT-2005-023253; SBL), Flanders Fund for Scientific Research (1841313N, G.0354.06, G.0758.10; and postdoc grant; Leuven1&2), Hartford Hospital (Olin), John Merck Scholars Fund (Yale), Kyulan Family Foundation (Trinity), Michigan Institute for Clinical and Health Research (MICHR) Pre-doctoral Fellowship (UM1&2), National Children’s Hospital (AMNCH; Trinity), National Initiative for Brain and Cognition NIHC HCMI (056-13-014, 056-13-017; SBL), NWO (051.07.003, 452-04-305, 400-08-089; SBL) and Netherlands Brain Foundation (KS 2010(1)-29; SBL), NRSA Pre-doctoral Fellowship (F31DC010143; USM), Research Council of the University of Leuven (Leuven1&2), Stanford Institute for Neuro-Innovations & Translational Neurosciences (Stanford), Leon Levy Foundation (NYU), Meath Foundation, Adelaide and Meath Hospital (Trinity), Stavros Niarchos Foundation (NYU), UCLA Autism Center of Excellence (UCLA1&2), and University of Utah Multidisciplinary Research Seed Grant (USM).

